# Hybrid histidine kinase BinK represses *Vibrio fischeri* biofilm signaling at multiple developmental stages

**DOI:** 10.1101/2021.03.29.437627

**Authors:** Denise A. Ludvik, Katherine M. Bultman, Mark J. Mandel

## Abstract

The symbiosis between the Hawaiian bobtail squid, *Euprymna scolopes*, and its exclusive light-organ symbiont, *Vibrio fischeri*, provides a natural system in which to study host-microbe specificity and gene regulation during the establishment of a mutually-beneficial symbiosis. Colonization of the host relies on bacterial biofilm-like aggregation in the squid mucus field. Symbiotic biofilm formation is controlled by a two-component signaling (TCS) system consisting of regulators RscS-SypF-SypG, which together direct transcription of the Syp symbiotic polysaccharide. TCS systems are broadly important for bacteria to sense environmental cues and then direct changes in behavior. Previously, we identified hybrid histidine kinase BinK as a strong negative regulator of *V. fischeri* biofilm regulation, and here we further explore the function of BinK. To inhibit biofilm formation, BinK requires the predicted phosphorylation sites in both the histidine kinase (H362) and receiver (D794) domains. Furthermore, we show that RscS is not essential for host colonization when *binK* is deleted from strain ES114, and imaging of aggregate size revealed no benefit to the presence of RscS in a background lacking BinK. Strains lacking RscS still suffered in competition. Finally, we show that BinK functions to inhibit biofilm gene expression in the light organ crypts, providing evidence for biofilm gene regulation at later stages of host colonization. Overall, this study provides direct evidence for opposing activities of RscS and BinK and yields novel insights into biofilm regulation during the maturation of a beneficial symbiosis.

**IMPORTANCE:** Bacteria are often in a biofilm state, and transitions between planktonic and biofilm lifestyles are important for pathogenic, beneficial, and environmental microbes. The critical nature of biofilm formation during *Vibrio fischeri* colonization of the Hawaiian bobtail squid light organ provides an opportunity to study development of this process *in vivo* using a combination of genetic and imaging approaches. The current work refines the signaling circuitry of the biofilm pathway in *V. fischeri*, provides evidence that biofilm regulatory changes occur in the host, and identifies BinK as one of the regulators of that process. This study provides information about how bacteria regulate biofilm gene expression in an intact animal host.

## INTRODUCTION

Animals are host to microbial partners that perform essential functions, including promoting tissue and immune development, nutrient acquisition, and defense (1, 2). In many cases, the hosts emerge aposymbiotic (i.e., lacking their symbiont) and must then recruit and retain the correct symbiotic microbes from the environment. On the microbial side, these horizontally-acquired symbionts often navigate multiple distinct lifestyles, including a free-living environmental stage and a distinct host-associated phase (3). To understand how reproducible symbiotic colonization occurs against this backdrop of distinct microbial life stages, we can use a model system in which a host organ is colonized by only a single bacterial partner. Here, we focus on marine *Vibrio fischeri* and their colonization of the light organ of the Hawaiian bobtail squid, *Euprymna scolopes* (4, 5). *V. fischeri* gains exclusive access to the squid’s light organ niche and creates luminescence that the squid manipulates as a counter-illumination camouflage strategy (6, 7). Within hours of the aposymbiotic squid hatching, *V. fischeri* colonize the light organ (8, 9). In addition to the binary nature of the symbiosis and the ability to rear both partners separately, the amenability of *V. fischeri* to sophisticated genetic manipulation and the power to image at the direct site of infection provides a powerful tool box to study how genes and their products influence bacterial colonization.

Biofilm formation by the symbiotic bacteria is fundamental to the colonization of the squid host (3, 10–15). During the establishment of the symbiosis, squid recruit *V. fischeri* by pumping seawater through the mantle cavity and over the light organ. Biofilm formation is required for *V. fischeri* to aggregate in the host mucus, and *V. fischeri* mutants unable to synthesize biofilm are unable to colonize the squid (10, 11). The polysaccharide component of the biofilm is the symbiosis polysaccharide (Syp), whose production is encoded by the 18-gene *syp* locus on the *V. fischeri* second chromosome (11). Expression of this locus is controlled by a two-component phosphorelay (**Figure 1A**). In strain ES114, the current model posits that hybrid sensor kinase RscS autophosphorylates at residue H412 in its dimerization and histidine phosphotransferase (DHp) domain and then relays the phosphoryl group to D709 in its receiver domain (REC) (16). The phosphoryl group is then relayed to residue H705 in the histidine phosphotransfer (HPt) domain of a distinct hybrid sensor kinase, SypF (17, 18). SypF then phosphorylates D53 in the REC domain of response regulator SypG (17, 19). Phospho-SypG acts as a σ^54^-dependent enhancer binding protein to promote transcription of the *syp* locus (11, 20). SypF has additional effects, including regulating the activity of serine kinase/phosphatase SypE to inactivate/activate its target SypA, which influences Syp biofilm formation downstream of *syp* transcription (21).

**Figure 1:**
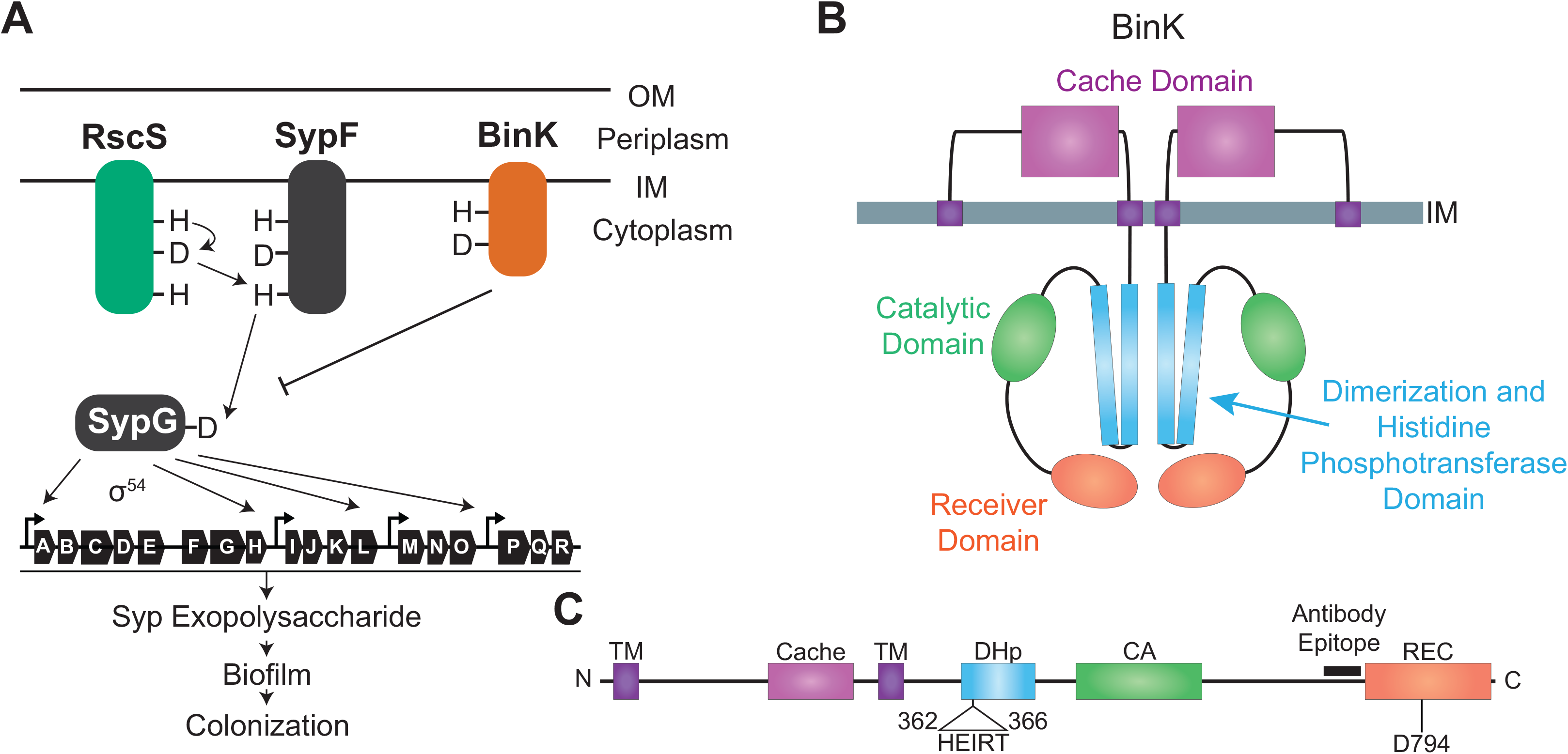
BinK signaling and domain organization. **A)** Model of the biofilm pathway in strain ES114 as is relevant for this study. RscS signals through SypF to SypG, which is a σ^54^-dependent transcriptional activator of the *syp* locus. Expression of the *syp* locus is required for biofilm formation. BinK inhibits biofilm production and feeds into the pathway at an unknown location at or above SypG. (OM, outer membrane; IM, inner membrane). **B)** Putative subcellular localization of a BinK homodimer **C)** BinK domain diagram. Positions of key residues and the epitope used to generate the BinK peptide antibody (722-742) are annotated. (TM, predicted transmembrane domains).

Previously, we identified the biofilm regulator, BinK, as a strong inhibitor of symbiotic biofilm formation (22). BinK was identified in an insertion sequencing screen where a mutant library of *V. fischeri* was analyzed before and after colonization of squid hatchlings to find genes that influenced colonization (23). Mutants of *binK* were overrepresented in the output pool compared to the input, suggesting that deletion of *binK* confers an advantage during the colonization process. Upon further study, it was determined that deletion of *binK* confers a competitive advantage against the wild-type strain during squid colonization, and the mutant forms larger aggregates on the surface of the squid light organ (22). Overexpressing BinK leads to a substantial reduction in symbiotic biofilm formation and prohibits bacteria from colonizing the host. BinK, which is conserved across *V. fischeri* strains, was also identified to be an important colonization regulator in a squid colonization experimental evolution study (14). Natural *V. fischeri* fish and seawater isolates that were experimentally evolved to colonize squid were able to do so as a result of spontaneous mutations in *binK* (14). Strain ES114 does not form robust biofilms outside of the squid. However, we can use culture-based assays with an RscS-overexpressing allele (termed, *rscS*\*) that approximates *in vivo* biofilm phenotypes to study BinK function, and with those approaches we demonstrated that deletion of *binK* can lead to wrinkled colony formation, higher transcription of the *syp* locus, and higher production of the Syp exopolysaccharide (22). Overexpressing *sypG* is epistatic to inhibitory signaling from overexpressing *binK* (22). The Δ*binK* strain is significantly derepressed for biofilm formation and in that background, calcium stimulates colony biofilm formation without the need for induction by *rscS*\* alleles (24). The calcium induction system led to the discovery of biofilm regulator HahK, which influences biofilm development in response to host nitric oxide (24, 25).

Given the prominence of BinK as a strong negative regulator of biofilm across various *V. fischeri* natural isolates, here we pursued multiple questions regarding its function during symbiosis. First, while BinK has the predicted structure of a hybrid histidine kinase, we asked whether it requires its putative phosphorylation sites for function. Second, given the strong phenotypes of BinK, we asked whether canonical squid isolate ES114 would be capable of colonizing squid in the absence of the positive regulator RscS if the negative regulator BinK was also removed. We found that RscS was dispensable for colonization in the absence of BinK, and this result enabled us to investigate the relative role of each protein during symbiotic colonization. Third, using direct imaging of a fluorescent *syp* transcriptional reporter, we asked whether symbiosis polysaccharide gene expression is regulated in the host by comparing regulation at two distinct time points (and therefore, distinct sites with the host) and by comparing both wild type and various mutant strains. Overall, this work provides an intriguing look into how biofilm signaling is regulated when a symbiotic microbe encounters its animal host.

## RESULTS

### BinK requires its conserved two-component histidine and aspartate for function

We identified BinK as an orphan hybrid histidine kinase that inhibits biofilm formation and is a negative regulator of squid colonization (22). Protein domain prediction indicated that BinK has the conserved dimerization and histidine phosphotransferase (DHp) domain and catalytic (CA) domain typical of a two-component sensor kinase, and it also contains an additional receiver (REC) domain making it a hybrid sensor kinase (**Figure 1B**) (26–29). His362 in the DHp domain and Asp794 in the REC domain are the predicted sites for phosphorylation in BinK (**Figure 1C**) (22). Phosphotransfer through such sites is typically required for signaling by sensor kinases, but there are examples where this is not the case (17, 30, 31). Therefore, we first asked whether His362 and Asp794 are necessary for BinK function.

To assess the function of individual alleles, we conducted colony biofilm assays. We started with a Δ*binK* strain and then introduced the *binK* gene (including 300 bp of upstream and downstream sequence) into the neutral chromosomal *att*Tn*7* site **(Figure 2A)**. This approach enabled us to test the wild-type and mutant alleles in comparable isogenic backgrounds. The strain background also has an allele to induce biofilm formation under culture conditions through the overexpression of RscS. This allele, termed *rscS*\* is carried on the chromosome at the native *rscS* locus (**Figure 2A**) (32). In this background, deletion of *binK* results in a wrinkled colony morphology when grown at 28 °C, while a strain with a functional *binK* has a smooth colony morphology (22). To test whether the His362 and/or Asp794 are required for BinK function, we constructed H362Q and D794A mutations, which have been shown to mimic the unphosphorylated state when similarly introduced into homologous domains (16, 33). In an otherwise Δ*binK* background, mutation in either individual residue or in both residues in the same protein resulted in a non-functional BinK (**Figure 2B**). We next asked whether a predicted phosphomimetic allele of the REC domain D794E is functional. This allele was constructed and unable to complement the lack of BinK in the *rscS*\* biofilm induction model (**Figure 2B**). Using western blot analysis with a polyclonal antibody raised against a BinK cytoplasmic epitope, we demonstrated that the mutant proteins are expressed to levels comparable to that of the wild-type BinK (**Figure 2C**). Together, these results provide genetic evidence that phosphorylation of BinK residues is required for its function. Given that we did not observe complementation with either the D794A that is predicted to be non-phosphorylatable or with the putative phosphomimetic D794E allele, we expect that phosphoryl groups at this residue are transferred to (or from) a downstream signaling partner, and that this signaling is required for BinK to inhibit biofilm formation.

**Figure 2:**
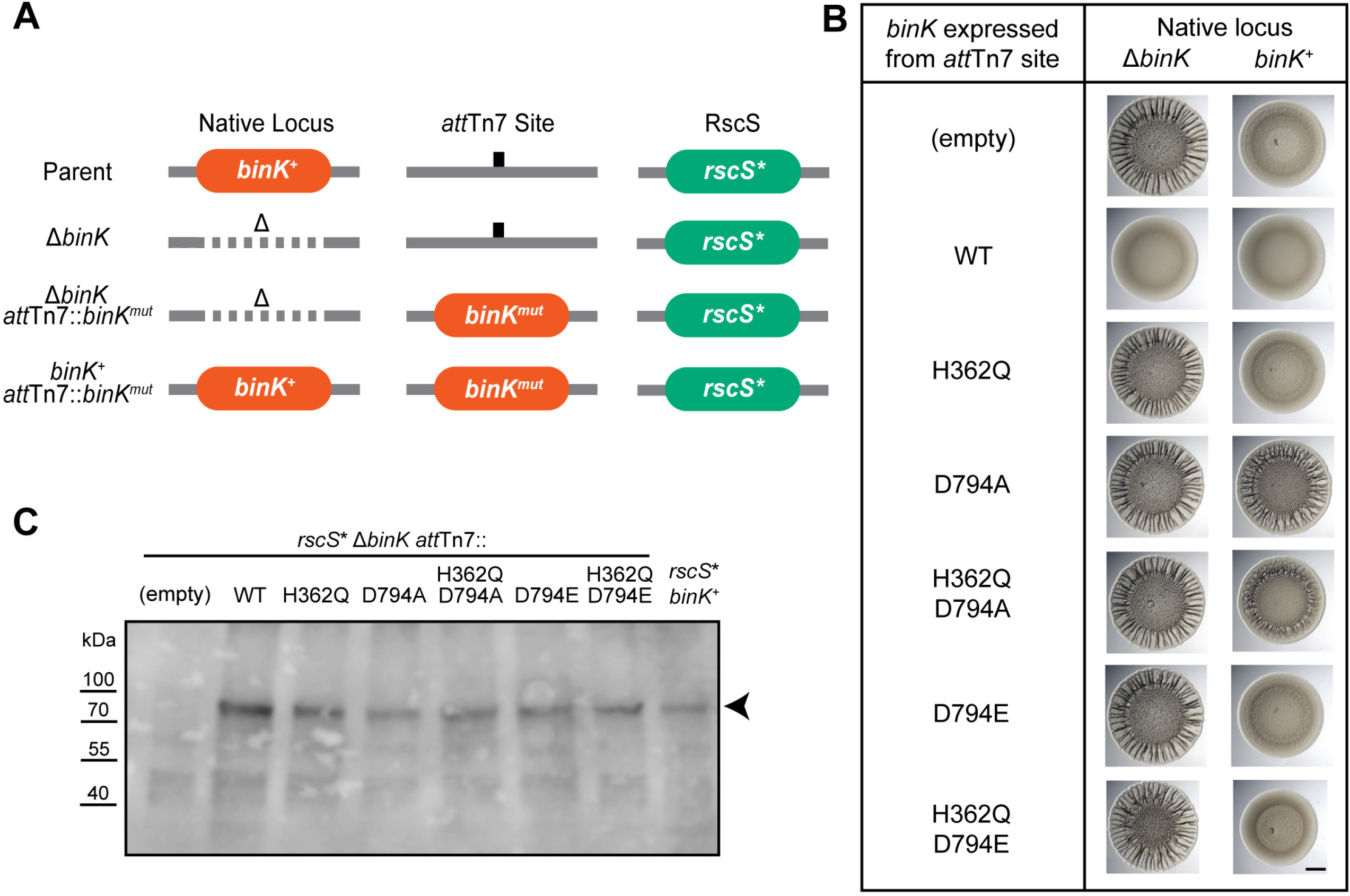
BinK requires H362 and D794 to inhibit colony biofilms. **A)** Genome representations of the different strains used to assess *binΚ* alleles. The parent ES114 *rscS*\* strain (MJM1198) was used to induce biofilm formation for wrinkled colony assays on plates. The *att*Tn*7* site is located on chromosome I and the native locus of *binK* is on chromosome II. *mut* designates a mutant allele of *binK*, while Δ indicates a clean deletion at that locus. **B)** Wrinkled colony assay of the strains indicated, grown at 28°C for 48 hours. Mutations (or WT control) expressed at the *att*Tn*7* site are listed beside each spot. In the left column, the expressed allele is the only *binK* allele in the cell, while in the right column wild-type *binK* is additionally present at its native locus. Scale bar is 2 mm. **C)** Western blot of whole cell lysates assessed with a peptide antibody against BinK. Arrow indicates BinK, which is predicted to be 97 kDa (61), and which is absent in the strain that does not encode the protein.

### BinK merodiploid analysis reveals a dominant negative phenotype for the D794A allele

Our data above demonstrated that the H362Q and D794A alleles were individually nonfunctional. Given that histidine kinases typically operate as homodimers, we next inquired whether the BinK(H362Q) or BinK(D794A) had any effect when expressed in a cell that also expressed wild-type BinK protein. We continued to express the test alleles from the *att*Tn*7* site, but now did so in a strain expressing wild-type *binK* from the native locus. In the wrinkled colony assay, the nonfunctional H362Q allele was recessive to the wild-type allele as expected, but the D794A allele exhibited a dominant negative phenotype, displaying a lack of biofilm inhibition even in the presence of the wild-type allele (**Figure 2B**). A similar dominance was observed for the D794A allele even if the H362Q mutation was introduced on the same polypeptide, expressed from the *att*Tn*7* site (**Figure 2B**). While either the H362Q or the D794A mutation disable BinK’s biofilm inhibitory function, the latter allele additionally interferes with the ability of a wild-type copy of BinK to inhibit biofilm formation.

Our data above demonstrated that the putative phosphomimetic D794E allele of BinK was similarly nonfunctional as the D794A allele when expressed as the only BinK allele in the cell. Merodiploid analysis revealed, however, that the D794E allele did not interfere with BinK activity *in trans* (**Figure 2B**). This was the case whether it was the only mutation in *binK* or in the context of a double *binK*(H362Q, D794E) allele (**Figure 2B**). Therefore, inactivation of the REC domain does not correlate directly with the dominant interfering phenotype. However, it may be the case that an unphosphorylated D794 (and the D794A allele that cannot be phosphorylated) leads to inhibition of BinK activity, whereas phosphorylated D794 (mimicked by the D794E allele) does not.

Sensor kinases can perform both phosphatase and kinase activities (34). To ask whether BinK kinase or phosphatase activity is used for biofilm inhibition, we created point mutations in the DHp domain that are predicted to differentially affect kinase versus phosphatase activities of the protein. BinK contains a conserved ExxT motif immediately after the conserved His362 residue (**Figure 1C**). This region of the H-box (i.e., the conserved phosphoryl group-binding His and surrounding region) is important for coordinating the phosphotransfer reactions in two-component proteins (35). In BinK we constructed E363A, T366Q, and T366A mutations that are predicted to eliminate kinase activity, reduce phosphatase activity, and eliminate phosphatase activity (with possible effects on autokinase activity), respectively (35–41). Colony biofilm assays revealed nonfunctional BinK in each case (**Figure S1**). This result further supports a role for phosphotransfer through BinK in its functional role.

### Single-copy phosphomimetic SypG is epistatic to BinK overexpression

As an orphan histidine kinase, BinK has no known paired response regulator. In a previous study, we demonstrated that a separate RscS-dependent pathway that relies on signaling through SypE was insensitive to BinK activity (22). Therefore, we can proceed to study BinK signaling architecture in a Δ*sypE sypF2* mutant background to isolate the core biofilm phosphorelay. Note that in this strain background, wrinkled colony formation is observed upon overexpression of SypG (without the requirement for *rscS*\*) (18, 42). In our previous study, we demonstrated that overexpression of SypG led the cell to be insensitive to overexpression of BinK in a wrinkled colony assay (22). This result suggested that SypG was epistatic to BinK. To further probe the genetic relationship between BinK and SypG, we constructed a phosphomimetic SypG allele at the native *sypG* locus. Hussa *et al.* demonstrated that a plasmid containing the SypG(D53E) allele conferred increased *syp* transcription and wrinkled colony formation in a Δ*sypE sypF2* strain (18). We introduced the same amino acid change into chromosomal *sypG* within the Δ*sypE sypF2* genetic background. We proceeded to determine that overexpression of BinK does not reduce the colony biofilm produced in the Δ*sypE sypF2 sypG*(D53E) background (**Figure S2**). Therefore, this experiment supports and extends our previous work and provides strong evidence that BinK acts upstream of SypG in the control of *syp* transcription and biofilm development. We proceeded then to examine the relative impacts of RscS and BinK on colonization phenotypes *in vivo*.

### RscS is not required for aggregation or squid colonization in a strain lacking BinK

In strain ES114, the positive regulator RscS and the negative regulator BinK both exert strong impacts on colonization and both act through the response regulator SypG. We therefore considered models in which RscS and BinK exert opposing influence on *syp* gene transcription during squid colonization. Deletion of *rscS* alone leads to severe *in vivo* biofilm and colonization defects, whereas deletion of *binK* alone leads to enhanced biofilm production and improved colonization in a competitive assay (10, 22, 43). Given these opposing phenotypes, we asked whether removal of both regulators--RscS and BinK--would enable the bacteria to colonize the host. An examination of experiments in diverse *V. fischeri* strains provides some insight into this question. Strains can improve their ability to colonize squid in the laboratory by mutation of *binK*, and this includes strain MJ11 that naturally lacks RscS (14, 22). The same MJ11 strain can colonize squid robustly if RscS from squid symbiont ES114 is introduced (13). Together, these results support our model that RscS and BinK could exhibit opposing activities, but we remained uncertain as to what we would observe in native squid symbiont ES114. If RscS is mainly required to counteract the negative regulation of BinK, then elimination of both regulators should allow the bacteria to colonize the squid host. However, if RscS is required to transduce a specific signal from the host, then we predict that elimination of both regulators would not allow for colonization.

To test these models, we conducted single-strain colonization assays. As has been shown previously, a Δ*rscS* mutant exhibits a significant colonization defect (**Figure 3**) (43). The Δ*binK* mutant is known to exhibit a competitive advantage over wild-type (22), yet in single-strain colonization displays similar bacterial yields and luminescence (**Figure 3**). The Δ*binK* Δ*rscS* double mutant strain was able to colonize up to levels indistinguishable from the wild-type strain (**Figure 3**). This result therefore supports the model that in ES114 BinK and RscS antagonize each others’ activity, and that in the absence of the negative regulator BinK, the positive regulator RscS is no longer required for single-strain colonization.

**Figure 3:**
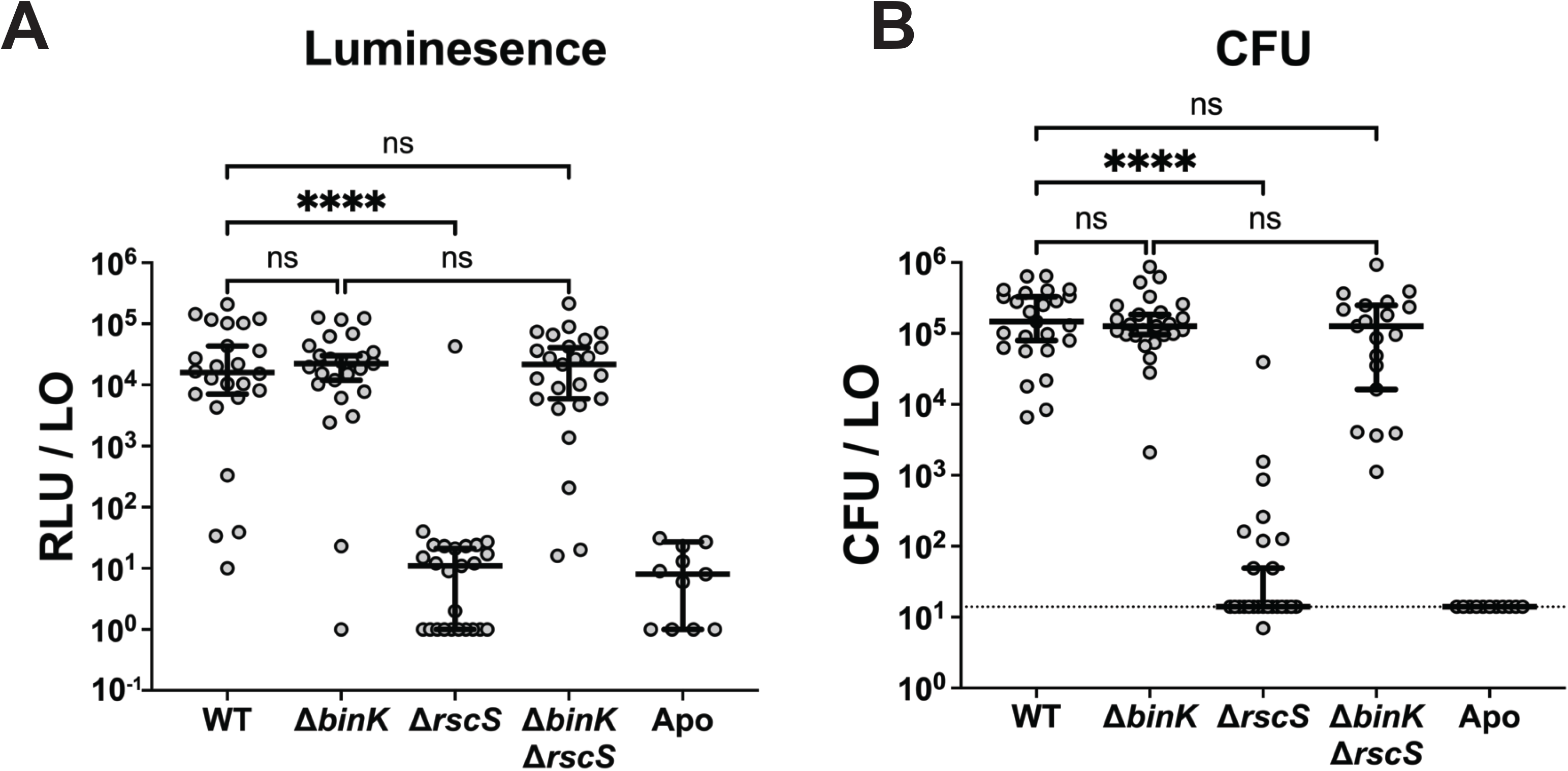
RscS is not required for squid colonization in a strain lacking BinK. Bacteria were inoculated into FSIO containing squid and allowed to colonize for 3 hours. Squid were washed and then maintained for two days to allow establishment of the symbiosis. Shown are data on the luminescence (A) and CFU (B) per light organ (LO) at 48 hpi. Each dot represents an individual animal. The dashed line indicates the limit of detection for CFU/LO. For both graphs, data is pooled from three replicate experiments. Bars represent the median for each strain with a 95% confidence interval. Statistical significance was calculated using Kruskal-Wallis test with Dunn’s multiple comparison tests. (ns, not significant; ****, p<0.0001)

The above result was surprising in that we identified a condition in which RscS, discovered twenty years ago as a strong colonization factor (43), was no longer required for squid colonization in strain ES114. This prompted us to ask whether the key symbiotic behavior regulated by RscS--*in vivo* aggregate formation--occurs in the Δ*binK* Δ*rscS* background. For this experiment, we introduced a plasmid that constitutively expresses GFP into the colonizing strains from Figure 3, and we asked whether these strains form biofilm aggregates in the squid mucus field. Direct visualization of the bacterial cells revealed the presence of biofilm aggregates in the host for the Δ*binK* Δ*rscS* cells (**Figure 4A**). Notably, the size of these aggregates were comparable to those formed by Δ*binK* single mutant cells, which were larger than the aggregates formed by wild-type *V. fischeri* (**Figure 4A,B**). This result therefore reveals that RscS is not required for aggregate formation in a background lacking BinK, and RscS does not contribute to aggregate size in this background.

**Figure 4:**
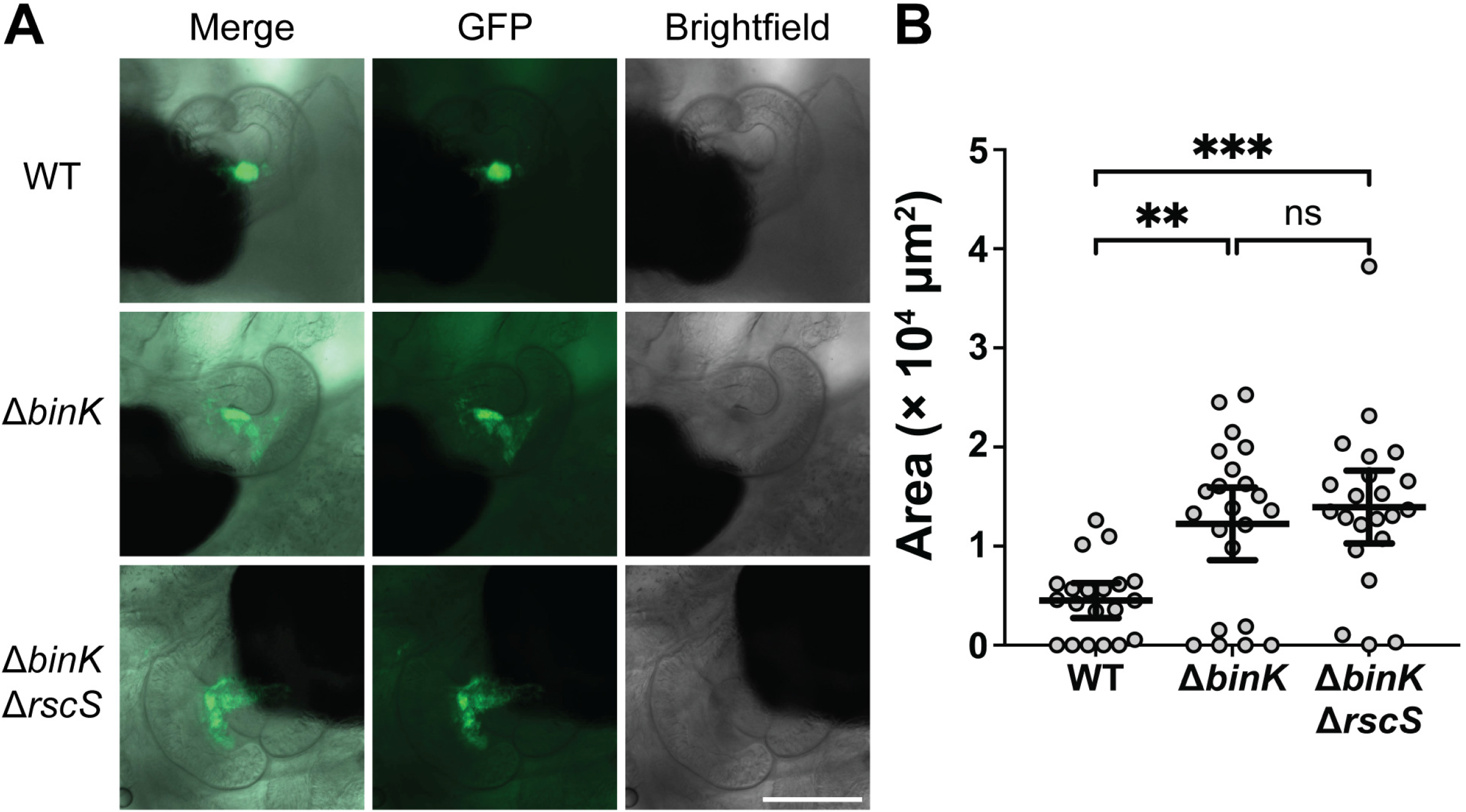
RscS does not impact aggregation in a strain lacking BinK. Imaging of bacterial aggregates in host mucus using fluorescence microscopy with the Zeiss Axio Zoom fluorescent microscope. Squid were inoculated with *V. fischeri* cells that constitutively express GFP from the pVSV102 plasmid and imaged at 3-4 hpi. **A)** Representative images of aggregates of approximately median size for each strain. Scale bar is 200 μm with all panels at the same scale. **B)** Quantification of aggregate area. Each dot represents one aggregate. A measure of zero indicates no aggregate was present. Significance was determined with a Kruskal-Wallis test and Dunn’s multiple comparison tests (ns, not significant; **, p<0.01; ***, p<0.001). The median area with a 95% confidence interval is displayed for each group. The data is pooled from two replicate experiments. WT = wild-type ES114 (MJM1100).

The above results prompted us to ask whether RscS performs any detectable function in a strain lacking BinK. We employed a sensitive competition assay to ask whether strains lacking RscS exhibit a defect upon co-inoculation. In a competitive colonization assay in which the Δ*binK* Δ*rscS* strain was co-inoculated with a LacZ-expressing Δ*binK* single mutant, the strain lacking RscS exhibited an approximately 100-fold defect in the competition (**Figure 5**).

**Figure 5:**
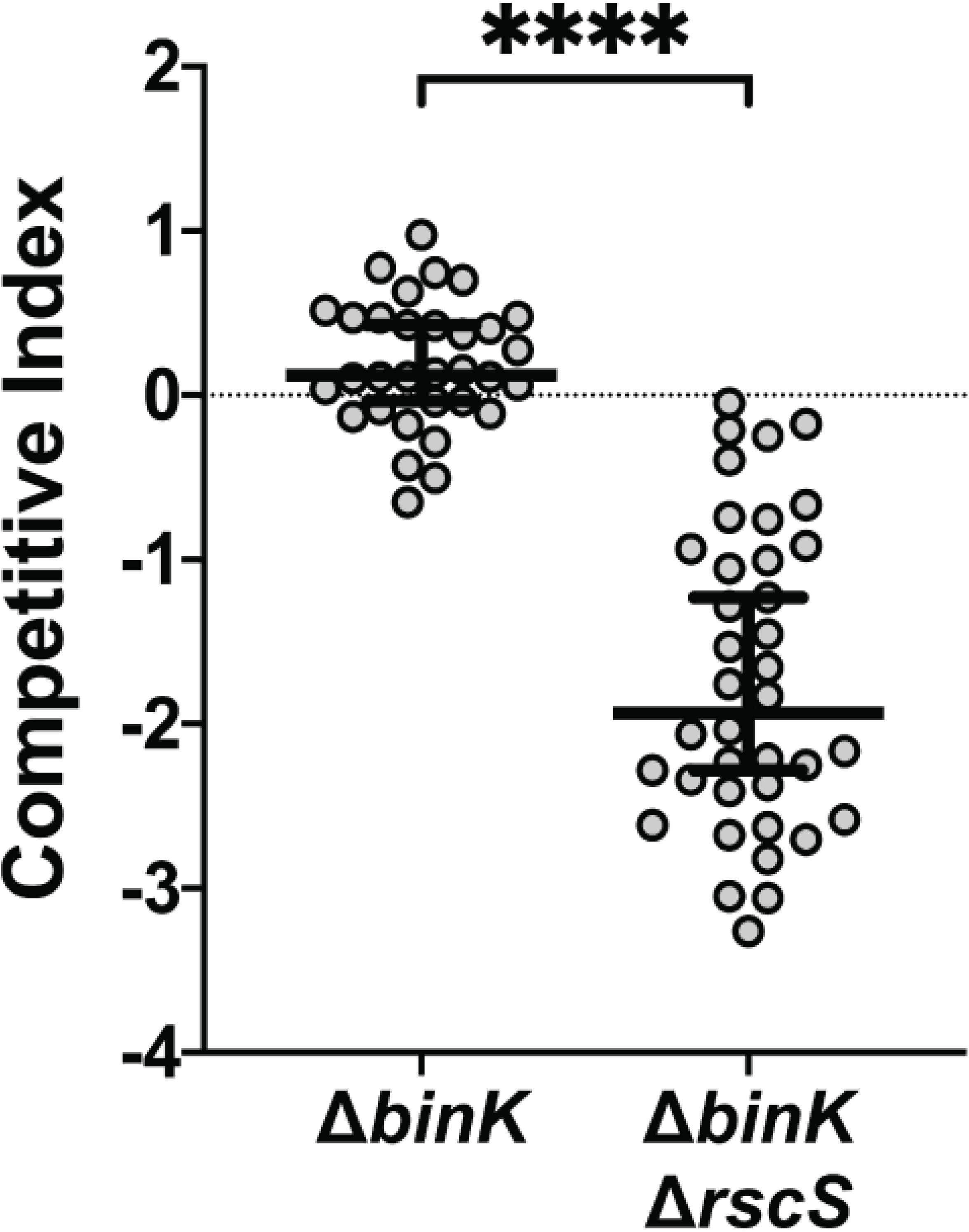
RscS is required for competitive fitness in a Δ*binK* background. Competitive fitness of the indicated strains (unlabeled) compared to a Δ*binK* strain (labeled with LacZ). Squid were exposed to a mixed inoculum of the two strains for 3 hours, then assessed at 48 hpi. Blue vs white CFU counts on LBS-Xgal were used to determine the representation of each strain in the competition. The competitive index is equal to the Log_10_-transformed value of the ratio (indicated strain/Δ*binK* strain) after competition normalized to its ratio in the input inoculum. Each point represents the competitive index from an individual squid. The median ratio with a 95% confidence interval is represented by the bar. Statistical significance was calculated with a Mann-Whitney U-test (****, p<0.0001).

### In the absence of BinK, *syp* transcription *in vitro* does not require RscS

Recent work demonstrated that in a strain lacking BinK, RscS is not required for colony biofilm formation when the symbiotic biofilm is induced with an additional 10 mM calcium in the medium (LBS-Ca) (24). We therefore examined a *sypA*’-*gfp*^+^ transcriptional reporter for cells grown on LBS-Ca agar, in strains that lack BinK, RscS, or both regulators. We determined that GFP activity from the reporter is induced in the Δ*binK* background, and the presence or absence of *rscS* in this background did not impact *syp* expression (**Figure 6A,C**). Similarly, the wrinkled colony biofilm phenotype is observed in both the Δ*binK* and Δ*binK* Δ*rscS* strains. Next, we examined the same reporter when the cells were grown on rich medium without the additional calcium (LBS). On LBS agar, the overall induction is lower than on LBS-Ca and strains do not wrinkle. However, we still detect induction of the reporter in the Δ*binK* strain compared to the WT parent. The induction is unaffected by the absence of *rscS* from this background (**Figure 6B,C**). In summary, we can readily detect *syp* gene transcription in the Δ*binK* background when grown on different solid media, and this induction is independent of RscS.

**Figure 6:**
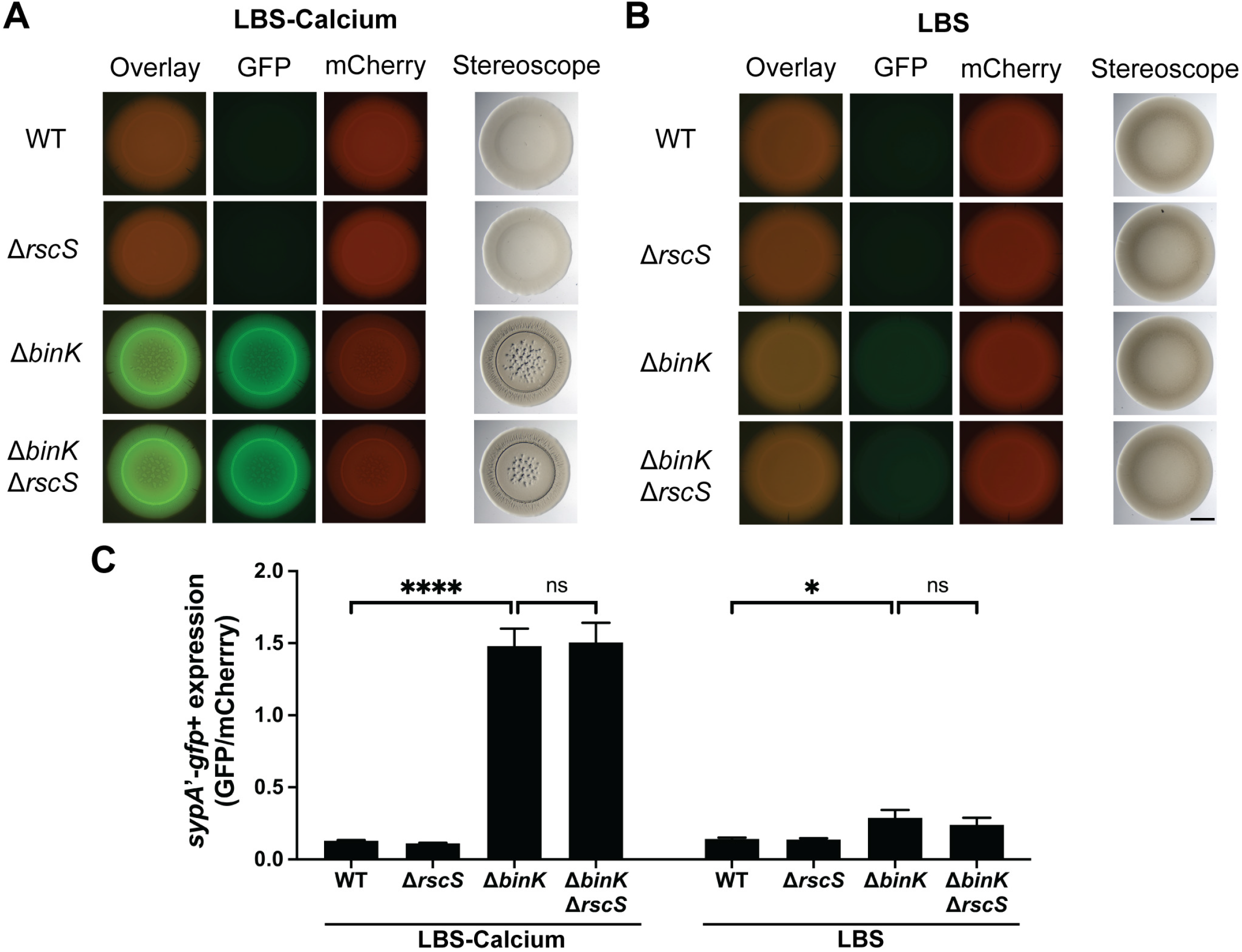
RscS does not alter *syp* production on agar in strains lacking BinK. *syp’-gfp*^+^ transcriptional reporter activity normalized to constitutive mCherry from strains carrying pM1422. **A, B)** Each strain was spotted onto LBS or LBS-Calcium and grown for 48 hours at 25°C. Images were taken on both a Zeiss Axio Zoom fluorescent microscope and Leica brightfield stereoscope. Scale bar in the bottom right of B is 2mm and all panels in A and B are at the same scale. **C)** Quantification of the fluorescence intensity. GFP intensity of the entire colony was normalized to mCherry intensity. Each bar represents the average GFP/mCherry of five different colonies from replicate experiments on two distinct days. Error bars are standard deviations. Statistical significance was determined with a two-way ANOVA analyzing the effect of strain background and media on fluorescence with Tukey’s test for multiple comparisons. (ns, not significant; *, p<0.05; ****, p<0.0001)

### BinK represses syp transcription in the squid crypts

We next examined expression of the *sypA*’-*gfp*^+^ transcriptional reporter in *V. fischeri* cells that had aggregated in the host mucus (3-4 hpi). The results in **Figure 7A,C** reveal indistinguishable overall levels of GFP activity in WT, Δ*binK*, and Δ*binK* Δ*rscS* cells in the aggregates of each animal examined. We note some limitations to these data. Given that cells spend a short period of time in the aggregate stage--on the order of 1-2 h--it is unclear whether the reporter is revealing the steady state transcription levels from the aggregate or whether this information integrates time prior to the aggregation stage (growth in liquid media and in seawater). Nonetheless, we present these data for two reasons. First, any physiological response that uses transcription to regulate symbiosis would be subject to similar constraints. Second, the similarity of the data points provides a useful control for the data in the crypts that will be described below. Even with these caveats, we can conclude that the absence of RscS does not diminish the median level of *sypA* transcription in the Δ*binK* background. We note that there was more heterogeneous *sypA’-gfp*^+^ activity across the aggregate in wild type compared to samples that lacked BinK (**Figure 7A**).

**Figure 7:**
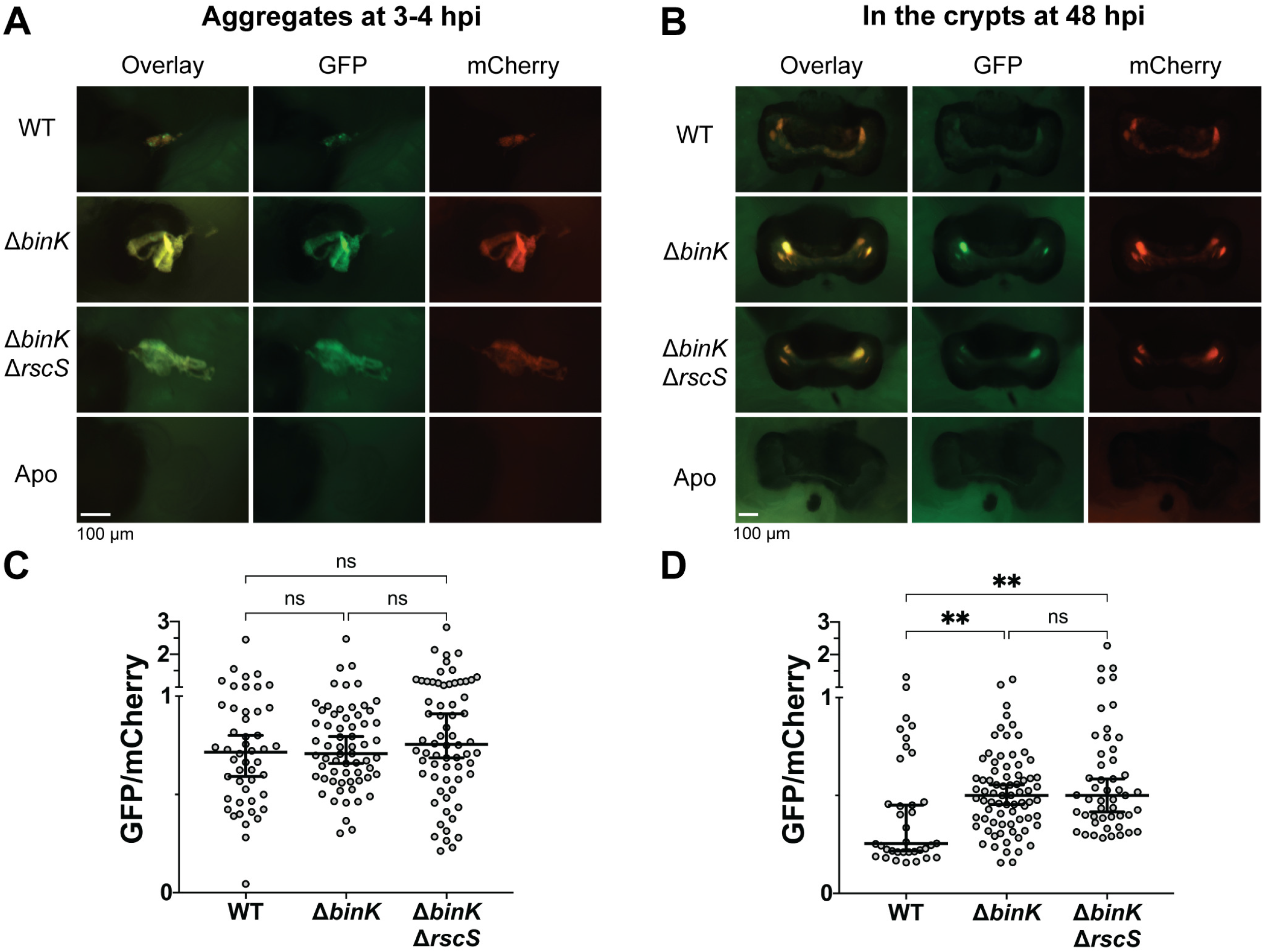
BinK inhibits *sypA* transcription in the crypts. Imaging of *V. fischeri* containing the pM1422 *sypA’-gfp*^+^ reporter plasmid (that encodes constitutive mCherry) while aggregating in host mucus at 3-4 hpi (A, quantified in C) or in host crypts at 48 hpi (B, quantified in D). For quantification, intensity of GFP and mCherry was measured for each individual aggregate or crypt, background signal was subtracted, and GFP was normalized to the mCherry level. Zen Blue software was used to collect signal and background measurements. Each dot represents an individual aggregate or crypt. Bars represent the median GFP/mCherry ratio with 95% confidence intervals. Statistical significance within each location was determined by a Kruskal-Wallis test with Dunn’s multiple comparison tests (ns, not significant; **, p<0.01). Data for each bacterial location is pooled from at least 3 replicate experiments that each contained 4-8 squid per strain.

We proceeded to conduct a similar analysis of the transcriptional reporter in the light organ crypts (48 hpi). At this point, we observed a notable difference between the Δ*binK* strain and wild type, with substantially elevated *sypA* transcription in the cells lacking BinK (**Figure 7B,D**). The absence of RscS did not affect the *sypA* reporter. From this imaging, we conclude that a normal function of BinK is to repress *syp* transcription in the crypts.

## DISCUSSION

By studying the *V. fischeri*-squid symbiosis model, we have refined our understanding of how bacterial biofilm signaling is regulated at the initiation of bacterial colonization. This study provides evidence that BinK acts as a hybrid histidine kinase, defines novel biofilm regulation in the host, describes a role for BinK in that regulation, and reveals that the key ES114 colonization factor RscS is dispensable in the absence of BinK. These major conclusions are discussed in detail below.

### BinK acts as a hybrid histidine kinase

In our previous study, we identified BinK as a putative hybrid histidine kinase based on its predicted domain structure containing CA, DHp, and REC domains (**Figure 1**) (22). Furthermore, experimental evolution studies to improve squid colonization of other *V. fischeri* strains revealed spontaneous *binK* point mutations in its CA and HAMP domains that approximated the phenotypes of Δ*binK* strains, suggesting key roles for these domains in BinK function (14). In this work, we used a combination of targeted mutants with a newly-developed anti-BinK peptide antibody to demonstrate that mutagenesis of predicted phosphorylation sites creates BinK proteins that are nonfunctional.

In most cases, histidine kinases dimerize, and our results provide genetic support that this occurs in the case of BinK. In particular, our finding that a BinK protein containing a REC domain that is locked as non-phosphorylated with the D794A mutation is not only nonfunctional, but is dominant negative as it interferes with signaling of a wild-type *binK* allele expressed in the same cell. There are published cases in which one monomer’s REC domain interacts with the same monomer’s DHp domain (*in cis*), and other cases in which it interacts with the partner monomer’s DHp domain (*in trans*) (44, 45). In both cases, however, dynamic movement of both monomers is required across the dimer interface (35, 46). It seems likely that the BinK D794A allele disrupts some aspect of this dynamic process that is still intact in heterodimers containing the phosphomimetic D794E allele. Therefore, BinK REC domain phosphorylation may both contribute to phosphoryl group flow and regulate the ability of BinK to interact in productive dimers.

Our data support a model in which phosphoryl groups flow from/to a downstream signaling partner through the BinK REC domain. The lack of complementation of a Δ*binK* allele with either the *binK*(D794A) or *binK*(D794E) alleles provide support for this model, in spite of differences in the merodiploid analysis described above. Additionally, the double *binK*(H362Q, D794E) allele is nonfunctional but not disruptive in the presence of a wild-type allele, suggesting that phosphoryl group flow can proceed *in cis* through a single BinK monomer even when present in mixed dimers with the double mutant. We note that there are examples in the literature where a REC domain regulates activity of the DHp domain and does not directly transfer phosphoryl groups (30, 31, 47). While this is possible for BinK, it seems unlikely given the genetic results here. Future biochemical analysis of BinK phosphotransfer will be necessary to further advance these studies. In particular, such analysis may enable asking whether BinK acts as a kinase or phosphatase when it inhibits biofilm formation. Without knowledge of the direct downstream partner of BinK, this question is difficult to access at present.

Considering that a phosphomimetic SypG is epistatic to BinK, it is likely that BinK is acting to control the phosphorylation of SypG. Our results indicate that the BinK His and Asp residues are necessary for activity. Thus, we predict BinK acts indirectly on SypG through the HPt domain in another regulator. Among known players, the HPt domain that has been shown to mediate phosphorylation of SypG is that of SypF (17, 24), making SypF a likely candidate. It is also possible that the relevant partner has yet to be identified. Genetic approaches to identify and characterize relevant partners will provide insight into the pathway downstream of BinK.

### RscS is dispensable for colonization in strain ES114 lacking BinK

We found that in strain ES114, *rscS* is not required for squid colonization in a Δ*binK* background. Strains lacking both regulators colonize to a level comparable to that of wild type, though we found through competitive colonization analysis that there still remains a role for RscS in this background **(Figures 3**, **5)**. There are three major phylogenetic groups of *V. fischeri* (15). Relevant for this work, the ancestral Group C strains include Mediteranean squid symbionts such as strain SR5. These strains encode functional BinK, but do not encode RscS, and it is unclear how they can colonize squid without the biofilm-promoting activity from RscS (15). Group C strains also include fish symbionts such as MJ11, which cannot colonize squid unless they gain RscS or lose BinK (13, 14). Derived from this group is Group B, which includes strain ES114, which is the focus of the present study. Squid symbionts in Group B typically encode both RscS and BinK, and mutation of RscS leads to an inability to colonize the squid (13, 15). In this study, it was found that when *binK* is deleted, strain ES114 no longer requires *rscS* for colonization (**Figure 3**). This result mirrors a previous finding in Group C, which found that strains lacking RscS (due to evolution) and also lacking BinK (due to directed mutation) could colonize squid (14). Given the diversity of biofilm regulation across *V. fischeri*, this result was not expected and highlights conserved aspects of regulation that are shared across much of the species (15). It therefore seems that one of the main functions of RscS is to antagonize BinK’s negative regulation of biofilm: without the negative regulator, the positive regulator is no longer absolutely required for host colonization. From the evolutionary tree, we know that *binK*--which is found throughout the species--predates the horizontal gene transfer event that enabled acquisition of *rscS*, which is only present in a derived group of *V. fischeri* (13, 15). Therefore, our work raises the question of whether there are other factors that antagonize BinK activity in strains that colonize squid independent of RscS (e.g., the Group C Mediterranean squid symbionts). It seems likely that such activity would be sufficient to enable colonization, given that mutations in *binK* facilitate colonization by strains that are otherwise unable to colonize well (14).

### BinK is a key regulator across the symbiotic life cycle

In a previous study, we used a fluorescent biofilm gene promoter fusion to examine gene expression in liquid medium (22). In this work, we expanded on that approach to examine expression of *sypA’-gfp*^+^ on solid medium and *in vivo* during colonization. On solid agar, we observed high levels of reporter expression under conditions known to induce Syp biofilm formation (LBS-Calcium (24)). We observed a less dramatic yet significant induction on medium where the Syp biofilm is not visibly apparent (LBS), revealing the sensitivity of this reporter. Our results provide evidence for expression of the *syp* biofilm in the aggregates, when biofilm formation is known to be required for colonization. We also demonstrate that in the absence of BinK, there is expression of *syp* reporter in the crypts, which supports a role for BinK in repressing biofilm gene expression at this later stage in the wild type strain. Building on our observation above that RscS is dispensable for colonization in strains lacking BinK, we observed similar levels of *syp* reporter expression in Δ*binK* and Δ*binK* Δ*rscS* strains. Our discovery that BinK functions to repress *syp* expression in the crypts hints that BinK regulation may serve a role in the daily expulsion of bacteria from squid at dawn. Little is known about how this process is regulated, yet it occurs daily in the mature symbiosis for the duration of the host’s lifetime. If this proves correct, it may help to explain why *binK* genes are widely conserved in *V. fischeri* despite the mutant having a competitive advantage (14, 15, 22). It seems likely that while the absence of BinK is beneficial to enter the squid host, the absence of the regulator (and subsequent inappropriate biofilm formation at later stages) may be detrimental to the daily homeostasis that is maintained long-term in the squid. We also examined *sypA’-gfp*^+^ reporter activity in biofilm aggregates. The average expression level within aggregates was similar for wild type or Δ*binK* mutant cells, but we observed greater heterogeneity in the expression in wild type. This suggests that in the presence of BinK, there is more variability in biofilm expression, and this is worthy of further study.

This work provides an exciting view into how biofilm gene expression is regulated *in vivo*. We know that within a few hours, planktonic bacteria transition to a biofilm state in the host mucus (3, 48). Despite a number of biofilm regulators being identified, how this process is controlled at the host interface is not well understood. Our results provide evidence that this regulation is dynamic over the course of colonization, as evidenced by the BinK-dependent repression that occurs specifically in the crypts at 48 hpi but is not evident in the aggregates at 3-4 hpi. We propose that the interaction of BinK with host-derived compounds may lead to downregulation of biofilm genes as bacteria transition from the biofilm aggregates during initiation to cells in the crypts during the persistence stage. Finally, we observed similar sizes of *in vivo* biofilm aggregates in the host in Δ*binK* and Δ*binK* Δ*rscS* strains, arguing that there is no requirement for stimulation of biofilm through RscS to initiate a productive symbiosis. We therefore posit that a key regulatory mechanism to control the planktonic-to-biofilm transition is host inhibition of BinK.

In summary, this work provides novel insight into the function of hybrid histidine kinase BinK, its relationship to RscS is regulating symbiotic biofilm formation, and the temporal control of symbiotic biofilm gene expression. Future work will continue to examine the signaling architecture downstream of BinK and host-derived molecules that may regulate BinK activity.

## MATERIALS AND METHODS

### Bacterial Strains, Plasmids, and Media

*V. fischeri* and *Escherichia coli* strains used in this study are listed in **Table 1**. Plasmids used in this study are listed in **Table 2**. *V. fischeri* strains were grown at 25°C or 28°C in Luria-Bertani salt (LBS) medium (25 g Difco LB broth [BD], 10 g NaCl, and 50 mL 1 M Tris buffer, pH 7.5, per liter). *E. coli* strains, used for cloning and conjugation, were grown shaking at 37°C in Luria-Bertani (LB) medium (25 g Difco LB broth [BD] per liter). Growth media were solidified with 1.5% agar (15 g Bacto agar (BD) per liter) as needed. When necessary, antibiotics were added to the media at the following concentrations: erythromycin, 5 μg/ml for *V. fischeri*; kanamycin, 100 μg/ml for *V. fischeri* and 50 μg/ml for *E. coli*; and chloramphenicol, 5 μg/ml for *V. fischeri* and 25 μg/ml for *E. coli*. The *E. coli* strain π3813 containing pKV496 is a thymidine auxotroph and was grown in LB with 50 μg/ml kanamycin supplemented with 0.3 mM thymidine (49, 50).

**Table 1:** Strains used in this study.

| Strain | Genotype | Source or Reference(s) |
| --- | --- | --- |
| ***V. fischeri*** |  |  |
| MJM1100 = ES114 | Natural isolate, squid light-organ | (56, 57) |
| MJM1107 | MJM1100 pVSV102 | (23) |
| MJM1198 | MJM1100 *rscS* * | (32) |
| MJM1438 | MJM1100 pM1422 | JF Brooks II |
| MJM1538 | MJM1100 pLostfoX-Kan | (23) |
| MJM1575 | MJM1100 pVSV103 | (22) |
| MJM2251 | MJM1100 $\Delta binK$ | (22) |
| MJM2252 | MJM2251 pLostfoX-Kan | (22) |
| MJM2255 | MJM1100 *rscS* * $\Delta binK$ | (22) |
| MJM2265 | MJM2251 pVSV102 | (22) |
| MJM2475 | MJM1198 *attTn7::binK* | (22) |
| MJM2476 | MJM2255 *attTn7::binK* | (22) |
| MJM2479 | MJM1198 *attTn7::erm* | (22) |
| MJM2480 | MJM2255 *attTn7::erm* | (22) |
| MJM2484 | MJM2255 *attTn7::binK*(H362Q) | JF Brooks II |
| MJM2493 | MJM2251 pM1422 | JF Brooks II |
| MJM2536 =<br>KV3299 | ES114 $\Delta sypE$ *sypF* * | (18, 42) |
| MJM3210 | MJM1198 *attTn7::binK*(H362Q) | This study |
| MJM3213 | MJM1198 *attTn7::binK*(D794A) | This study |
| MJM3215 | MJM2255 *attTn7::binK*(D794A) | This study |
| MJM3218 | MJM1198 *attTn7::binK*(H362Q,D794A) | This study |
| MJM3220 | MJM2255 *attTn7::binK*(H362Q,D794A) | This study |
| MJM3221 | MJM2251 pVSV103 | This study |
| MJM3236 | MJM2536 *sypG*(D53E) | This study |
| MJM3251 | MJM1198 *attTn7::binK*(D794E) | This study |
| MJM3252 | MJM2255 *attTn7::binK*(D794E) | This study |
| MJM3256 | MJM1198 *attTn7::binK*(E366A) | This study |
| MJM3257 | MJM2255 $\Delta binK$ *attTn7::binK*(E363A) | This study |
| MJM3775 | MJM1100 $\Delta rscS::erm$-bar | This study |
| MJM3903 | MJM1100 $\Delta rscS::bar$ | This study |
| MJM4017 | MJM1100 $\Delta binK \Delta rscS::erm$-bar | This study |
| MJM4018 | MJM1100 $\Delta binK \Delta rscS::bar$ | This study |
| MJM4071 | MJM4018 pVSV102 | This study |
| MJM4240 | MJM3236 pVSV104 | This study |
| MJM4241 | MJM3236 pBinK | This study |
| MJM4242 | KV3299 pVSV104 | This study |
| MJM4243 | KV3299 pBinK | This study |
| MJM4251 | MJM1198 *attTn7::binK*(H362Q,D794E) | This study |
| MJM4252 | MJM2255 *attTn7::binK*(H362Q,D794E) | This study |
| MJM4253 | MJM1198 *attTn7::binK*(T366Q) | This study |
| MJM4254 | MJM2255 $\Delta binK$ *attTn7::binK*(T366Q) | This study |
| MJM4255 | MJM1198 *attTn7::binK*(T366A) | This study |
| MJM4256 | MJM2255 $\Delta binK$ *attTn7::binK*(T366A) | This study |
| MJM4257 | MJM3903 pM1422 | This study |
| MJM4258 | MJM4018 pM1422 | This study |
| ***E. coli*** |  |  |
| MJM534 | CC118 λpir/pEVS104 | (51) |
| MJM537 | DH5α λpir | Laboratory Stock |
| MJM542 | DH5α λpir/pVSV102 | (58) |
| MJM570 | DH5α/pEVS79 | (51) |
| MJM580 | DH5α λpir/pVSV104 | (58) |
| MJM637 | S17-1 λpir/pUX-BF13 | (59) |
| MJM658 | DH5α λpir/pEVS107 | (60) |
| MJM1422 | DH5α λpir/pM1422 | (22) |
| MJM2090 | DH5α λpir/pLostfoX-Kan | (23) |
| MJM2384 | DH5α λpir/pBinK | (22) |
| MJM2474 | DH5α λpir/pTn7BinK | (22) |
| MJM2482 | DH5α λpir/pTn7BinK(H362Q) | JF Brooks II |
| MJM3211 | DH5α λpir/pDAT01 | This study |
| MJM3216 | DH5α λpir/pDAT02 | This study |
| MJM3234 | DH5α λpir/pDAT05 | This study |
| MJM3241 | DH5α λpir/pDAT10 | This study |
| MJM3255 | DH5α λpir/pDAT12 | This study |
| MJM3478 | π3813/pKV496 | (50) |
| MJM4248 | DH5α λpir/pDAT13 | This study |
| MJM4249 | DH5α λpir/pDAT14 | This study |
| MJM4250 | DH5α λpir/pDAT15 | This study |

**Table 2:** Plasmid List.

| Plasmid | Description | Source or Reference(s) |
| --- | --- | --- |
| pEVS107 | Mini-Tn7 mobilizable vector (Kan<sup>R</sup>,Erm<sup>R</sup>) | (60) |
| pTn7BinK | pEVS107 carrying wild-type *binK* (Kan<sup>R</sup>,Erm<sup>R</sup>) | (22) |
| pTn7BinK(H362Q) | pEVS107 carrying *binK* with H362Q mutation (Kan<sup>R</sup>,Erm<sup>R</sup>) | This study |
| pDAT01 | pEVS107 carrying *binK* with D794A mutation (Kan<sup>R</sup>,Erm<sup>R</sup>) | This study |
| pDAT02 | pEVS107 carrying *binK* with H362Q,D794A mutation (Kan<sup>R</sup>,Erm<sup>R</sup>) | This study |
| pDAT10 | pEVS107 carrying *binK* with D794E mutation (Kan<sup>R</sup>,Erm<sup>R</sup>) | This study |
| pDAT13 | pEVS107 carrying *binK* with H362Q,D794E mutation (Kan<sup>R</sup>,Erm<sup>R</sup>) | This study |
| pDAT12 | pEVS107 carrying *binK* with E363A mutation (Kan<sup>R</sup>,Erm<sup>R</sup>) | This study |
| pDAT14 | pEVS107 carrying *binK* with T366Q mutation (Kan<sup>R</sup>,Erm<sup>R</sup>) | This study |
| pDAT15 | pEVS107 carrying *binK* with T366A mutation (Kan<sup>R</sup>,Erm<sup>R</sup>) | This study |
| pEVS104 | Conjugal helper plasmid (Kan<sup>R</sup>) | (51) |
| pUX-BF13 | Tn7 transposition helper (Amp<sup>R</sup>) | (59) |
| pVSV104 | Vector backbone for complementation (Kan<sup>R</sup>) | (58) |
| pBinK | pVSV104 carrying wild-type *binK* (Kan<sup>R</sup>) | (22) |
| pEVS79 | Vector backbone for allelic exchange (Cam<sup>R</sup>) | (51) |
| pDAT05 | pEVS79 carrying *sypG* with D53E mutation (Cam<sup>R</sup>) | This study |
| pM1422 | pTM267 *sypA'*-gfp+ (Cam<sup>R</sup>) | (22) |
| pVSV102 | Constitutive GFP (Kan<sup>R</sup>) | (58) |
| pVSV103 | Constitutive LacZ (Kan<sup>R</sup>) | (58) |
| pLostfoX-Kan | Arabinose-inducible TfoX for transformation (Kan<sup>R</sup>) | (23) |
| pKV496 | pEVS79 containing the FLP recombinase (Kan<sup>R</sup>) | (50) |

### DNA synthesis and sequencing

Each of the primers listed in **Table 3** was synthesized by Integrated DNA Technologies (Coralville, IA). Full inserts from all cloned constructs were verified by Sanger DNA sequencing at Northwestern University Feinberg School of Medicine Center for Genetic Medicine, Functional Biosciences via UW-Madison, or the UW-Madison Biotechnology Center. Sequence data were analyzed with SeqMan Pro (DNAStar software) and Benchling. For cloning and sequencing PCR reactions, we used Q5 High-Fidelity DNA polymerase (NEB). For diagnostic PCR, we used GoTaq polymerase (Promega).

**Table 3:**
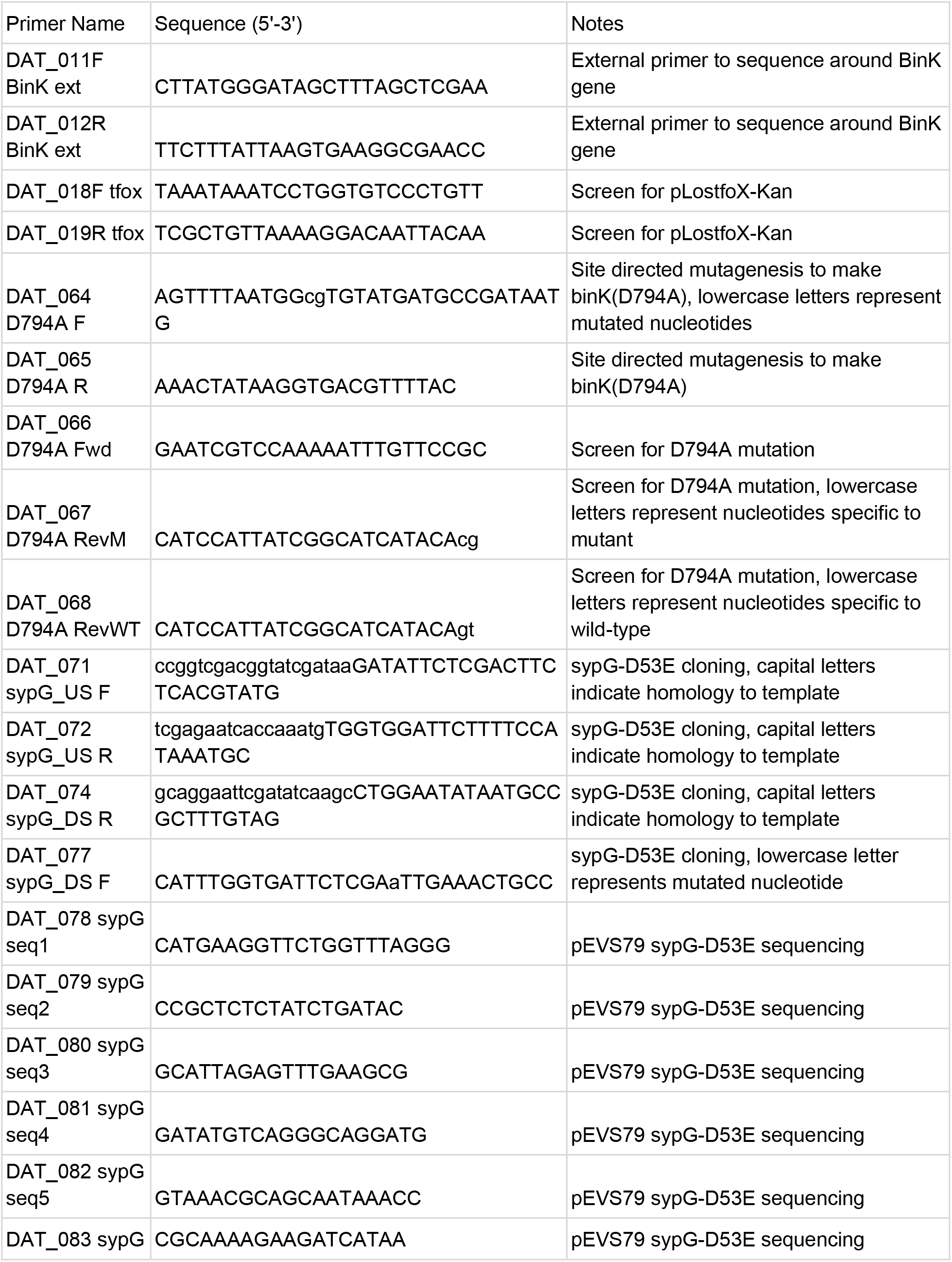
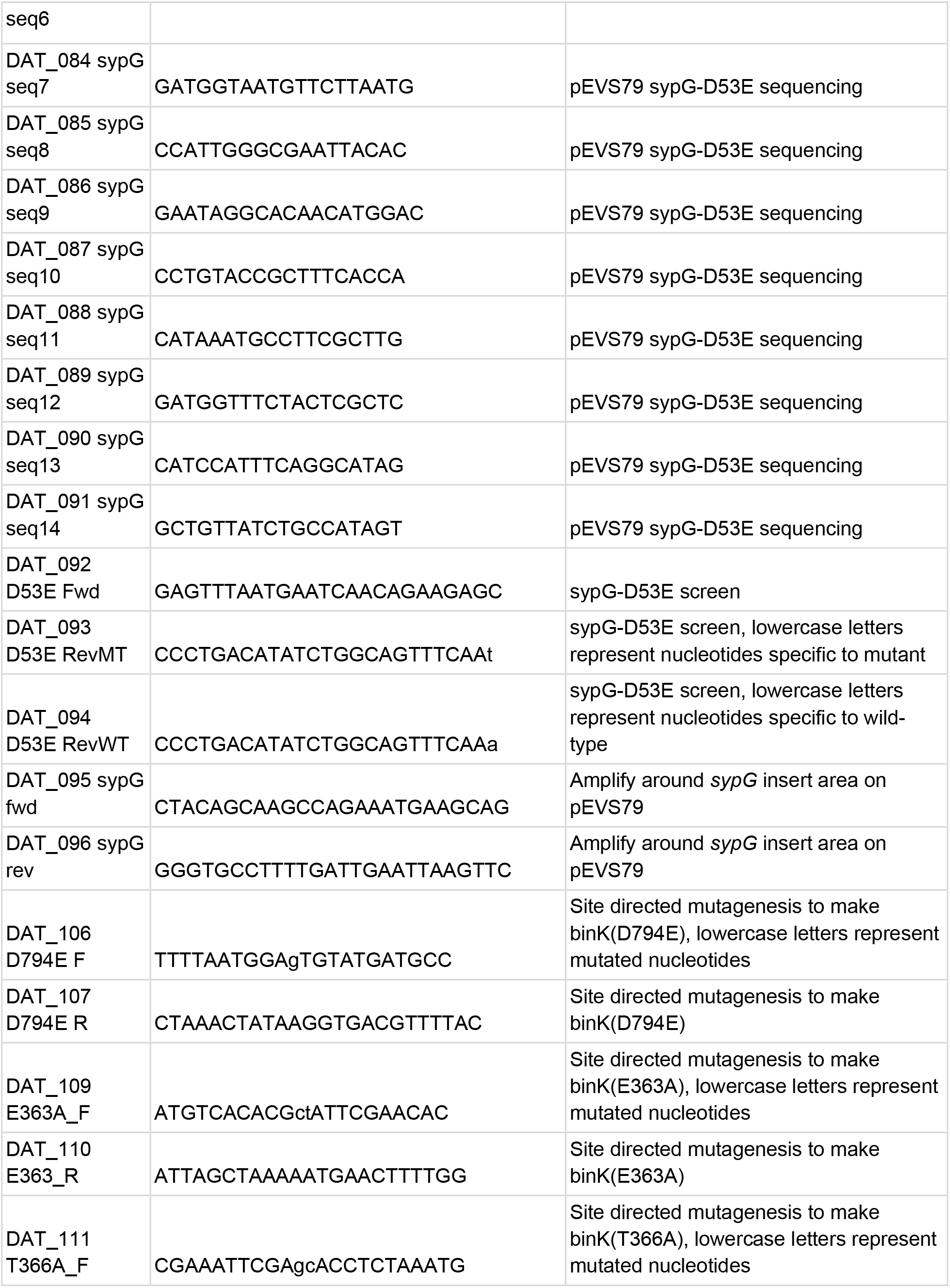
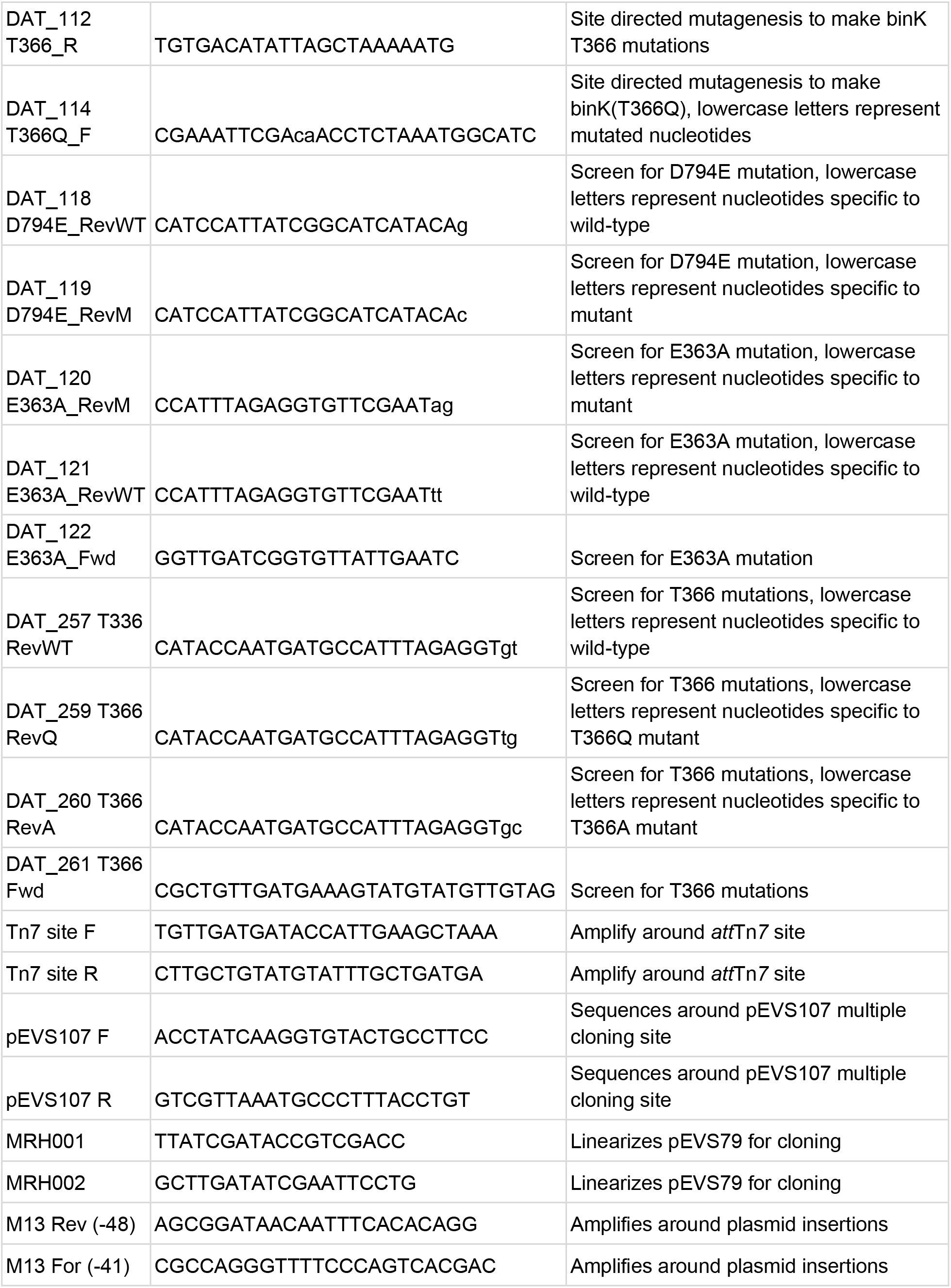
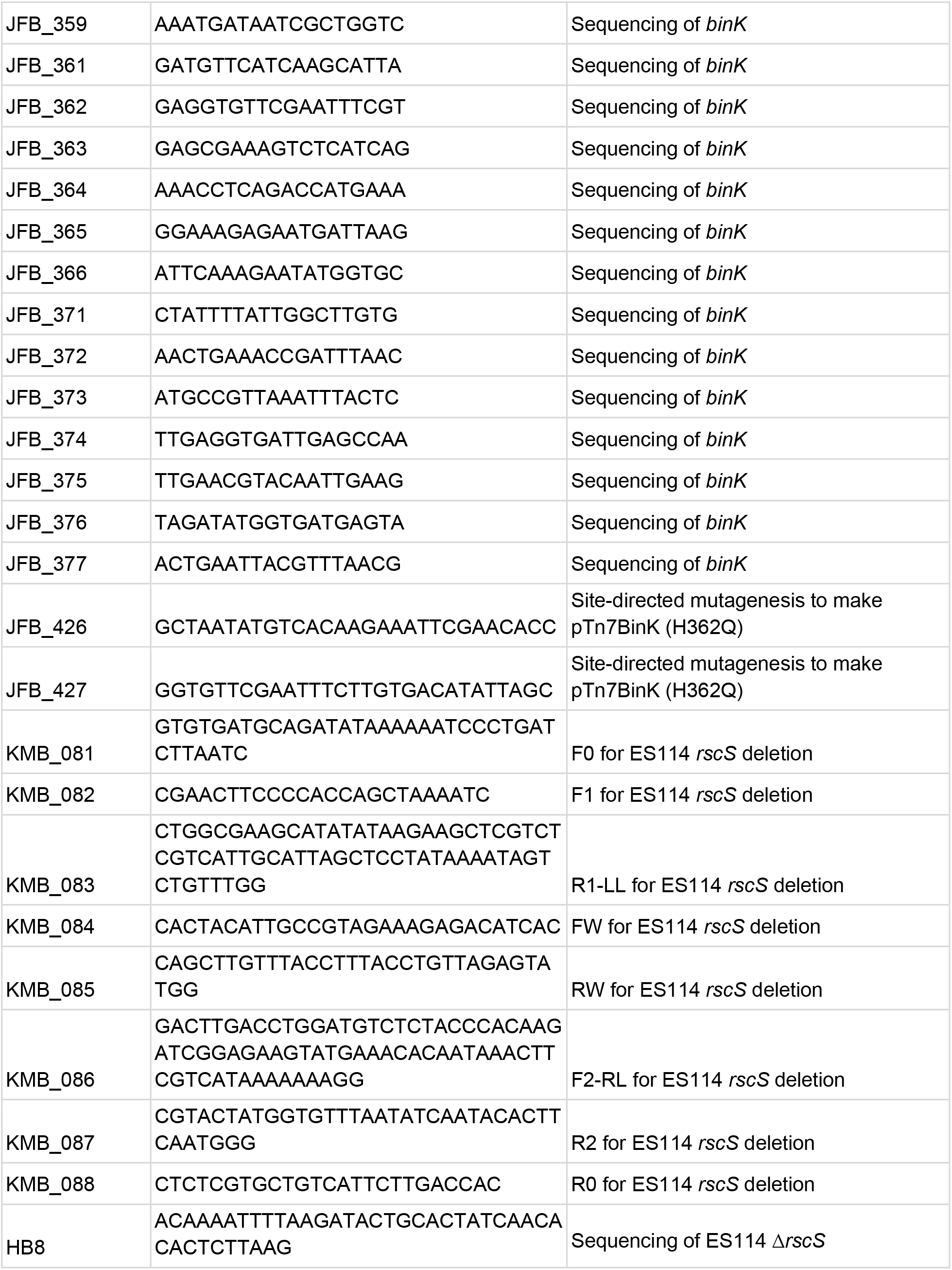
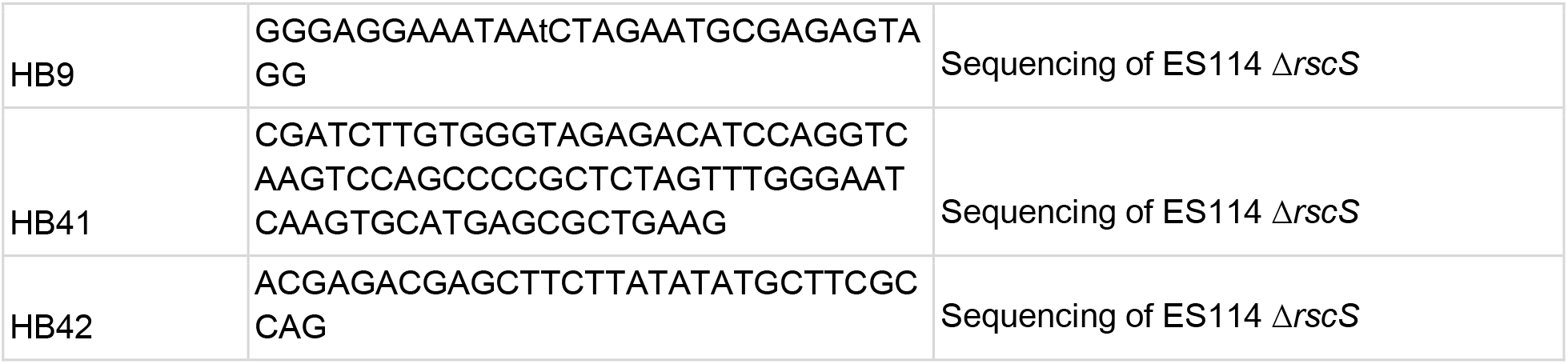
Primer List.

### Construction of*att*Tn*7*::*binK* mutant alleles

The previously generated pTn7-*binK* plasmid, which uses a mini-Tn*7* delivery vector backbone (pEVS107), was purified and used as a template. Point mutations to the *binK* sequence on the plasmid were designed using the NEBaseChanger tool and constructed with the Q5 site-directed mutagenesis kit (New England BioLabs, Inc.). The constructed plasmid was transformed into either electrocompetent or chemically competent DH5α λpir *E. coli*. The entire *binK* gene on the plasmid construct was sequenced (using pEVS107 F and R primers and *binK* sequencing primers). BinK alleles generated in this manner were then introduced into *V. fischeri* by tetraparental mating by mixing the pEVS104-containing helper, pUX-BF13-containing transposase, pEVS107 mini-Tn*7* vector containing donor, and the *V. fischeri* recipient (51). PCR verification by amplifying around the *att*Tn*7* site with primers Tn7 site F and Tn7 site R confirmed transposon insertion at the *att*Tn*7* site.

### Wrinkled Colony Assays

Cultures were grown overnight and 8 μL was spotted onto LBS plates or LBS-Calcium (10 mM CaCl_2_). Plates were incubated at 25°C or 28°C for 48 hours and imaged using Leica M60 stereomicroscope with Leica Firecam software. For assays done with the pM1422 reporter, plates were also imaged on a Zeiss Axio Zoom.v16 large-field fluorescent stereo microscope and analyzed with Zen Blue software.

### BinK peptide antibody creation and purification

ProSci (Poway, CA) analyzed the sequence of BinK and chose the peptide SEYGEMIDLPHKRKDNLDIIK as an epitope for polyclonal antibody production. This peptide was synthesized by ProSci and used to inoculate rabbits. Serum was analyzed at specific checkpoints to assess antibody production. Final bleed serum was then purified in our laboratory using a Proteus Protein A Mini Purification Starter Kit and yielded approximately 1 mg/mL of the antibody. Purified antibody was diluted 1:1 in 50% glycerol, aliquoted into small volumes, and stored at −20°C. Single aliquots were thawed and used for each blot.

### SDS-PAGE and Western Blots

One milliliter of overnight culture was pelleted, washed, and lysed in 1% SDS. The volume of SDS used to lyse the sample was adjusted based on the OD_600_ of the overnight culture to standardize the concentration of total protein in the samples. The solution was then pelleted to remove cell debris and the supernatant was mixed 1:1 with 2x Laemli sample buffer (Bio-Rad) and beta-mercaptoethanol. Samples were heated at 95°C for 15 minutes and loaded onto a 10% Bio-Rad Mini-PROTEAN TGX Precast Gel. The gel was then transferred to a PVDF membrane and blocked overnight in 5% non-fat milk. The purified anti-BinK-peptide antibody was used as the primary antibody in a 1:100 dilution in 0.5% nonfat milk in 1X TBS-Tween20. The secondary antibody was a 1:5,000 dilution of the Pierce goat anti-rabbit IgG (H+L) HRP-conjugate (Lot UK293475). Washes were done in 1X TBS-Tween20. Blots were developed using the ThermoScientific SuperSignal West Dura Extended Duration Substrate and analyzed using a Licor Odyssey Fc machine.

### Construction of *sypG*(D53E)

Site-direction mutation of *sypG* was done using an allelic exchange approach modified from the laboratory’s gene deletion protocol (https://doi.org/10.5281/zenodo.1470836). In brief, approximately 1.6-kb upstream sequence and 1.6-kb downstream sequence of the desired point mutation site in *sypG* was amplified from ES114 genomic DNA using primers designed using the NEB site-directed mutagenesis approach. These two fragments were then cloned into pEVS79 (which had been linearized by primers MRH001 and MRH002) using isothermal assembly (NEBuilder HiFi DNA assembly cloning kit) with the primer combinations listed in **Table 3**. The reaction was transformed into *E. coli* with selection for transformants on LB-chloramphenicol. PCR around the insertion using primers M13 For (−41) and M13 Rev (−48) was used to confirm plasmid candidates. The resulting candidates were then confirmed by sequencing and conjugated into the *V. fischeri* recipient (KV3299/MJM2536) by triparental mating with helper plasmid pEVS104, selecting for the chloramphenicol resistance of the plasmid backbone. Single recombinants in *V. fischeri* were screened for maintaining chloramphenicol resistance. To obtain double recombinants, single recombinants were then grown without antibiotics and patched onto LBS and LBS-Cam to find isolates that lost the antibiotic resistance cassette. These candidates were then verified with PCR and sequencing to confirm loss of the *cam* cassette and mutation of *sypG* to *sypG*(D53E). Strain MJM2536 *sypG*(D53E) was saved as MJM3236.

### Construction of Δ*rscS* and Δ*binK* Δ*rscS* strains

Deletion of *rscS* was performed following the barcode-tagged gene deletion protocol from Burgos et al. (52). In brief, the US homology arm was amplified using primers KMB_082 and KMB_083 and the DS homology arm was amplified using primers KMB_086 and KMB_087. Homology arms were fused to either side of a third fragment containing an *erm* cassette using SOE-PCR. Mutagenic DNA was purified using the Qiagen PCR purification kit and transformed into ES114 via transformation using pLostfoX-Kan (MJM1538) (53, 54). Mutant candidates were selected using erythromycin and screened by PCR using primer pairs KMB_081/KMB_088, KMB_081/HB8, and KMB_084/KMB_085. Insertion of the Erm-bar scar was confirmed by Sanger sequencing using primers KMB_081, KMB_082, KMB_083, HB8, HB9, KMB_086, KMB_087, KMB_088 and the barcode sequence recorded. The final bar-scar strain (MJM3903) was constructed via a triparental mating with donor MJM3478 (π3813/pKV496) (50) and helper strain MJM534 (CC118 λpir/pEVS104) with MJM3775. Candidates were selected for using kanamycin and screened by PCR using the primer pairs listed above. The deletion scar was verified by Sanger sequencing using primers KMB_081, KMB_082, KMB_083, HB41, HB42, KMB_086, KMB_087 and KMB_088.

To create the Δ*binK* Δ*rscS* strain, a Δ*binK* strain with the pLostfoX-Kan plasmid (MJM2252) was cultured for transformation (54). Donor DNA was 2.4 ug of genomic DNA from the Δ*rscS*::*erm-bar* strain that was purified using the Qiagen Blood and Tissue Kit. Transformed cells were plated onto LBS containing erythromycin to select for transformants. Isolates were then patched onto LBS containing kanamycin to ensure loss of the pLostfoX-Kan plasmid. PCR was performed to ensure the strain was transformed with Δ*rscS*::*erm-bar* and maintained Δ*binK* (primers DAT_011F BinK ext, DAT_012R BinK ext, KMB_081, KMB_088). The *erm* cassette was then removed using a FLP recombinase as described with the Δ*rscS::bar* strain above to create the final Δ*binK* Δ*rscS*::*bar* strain (MJM4018).

### Squid single strain colonizations

*V. fischeri* strains were grown overnight with aeration at 25°C in LBS. Overnight cultures were diluted 1:80 in LBS and grown to an OD_600_ of approximately 0.3. The OD_600_ was used to normalize the amount of each strain used to inoculate *E. scolopes* hatchlings at concentrations of approximately 6 × 10^3^ CFU/mL in 40 mL seawater for 3 hours. Squid were then washed and transferred to individual vials with 4 mL of bacteria-free filter-sterilized Instant Ocean (FSIO) until approximately 48 hours post-inoculation (hpi) with a water change that occurred at 24 hpi. At 48 hpi, squid were transferred to 1.5 mL microfuge tubes with 750 mL of water and each animal’s luminescence was measured using the Promega GloMax 20/20 luminometer.

CFU counts per light organ were conducted as we described previously: euthanized squid were homogenized and plated, and colonies were counted to determine CFU per light organ (55).

For crypt visualization, squid were anesthetized in FSIO with 2% ethanol. At this point, hatchlings were either immediately dissected and imaged, or fixed in 4% paraformaldehyde in 1X mPBS (50 mM phosphate buffer, 0.45 M NaCl, pH 7.4) for approximately 36 hours. Fixed squid were thoroughly washed in 1X mPBS before being dissected and imaged. All images were acquired on the Zeiss Axio Zoom.v16 large-field stereo microscope. The Zen Blue software polygon tool was used to select regions of interest and measure fluorescence intensity.

### Squid aggregation assays

*V. fischeri* strains were grown overnight with aeration at 25°C in LBS. 40 mL of overnight culture was used to inoculate *E. scolopes* hatchlings at concentrations of approximately 5.5 × 10^6^ CFU/mL in 40 mL for 3-4 hours. Hatchlings were then anesthetized in FSIO with 2% ethanol. At this point, hatchlings were either immediately dissected and imaged, or fixed in 4% paraformaldehyde in 1X mPBS (50 mM phosphate buffer, 0.45 M NaCl, pH 7.4) for approximately 36 hours. Fixed squid were thoroughly washed in 1X mPBS before being dissected and imaged. All images were acquired on the Zeiss Axio Zoom.v16 large-field fluorescent stereo microscope. The Zen Blue software polygon tool was used to select regions of interest and measure both area and fluorescence intensity

### Squid competition assays

Strains were grown overnight with aeration at 25°C in LBS and LBS containing kanamycin to maintain plasmid pVSV103. Strains with pVSV103 constitutively express LacZ (β-galactosidase). Overnight cultures were diluted 1:80 in LBS and grown to an OD_600_ of approximately 0.3. Using optical density to normalize the strains, the two strains were mixed in a 1:1 ratio. This mixed culture was used to inoculate *E. scolopes* hatchlings at concentrations of approximately 7.6 × 10^3^ bacteria for 3 hours. Squid were then washed and transferred to 40 ml of bacteria-free filter-sterilized Instant Ocean (FSIO) until approximately 48 hours post-inoculation (water was changed at 24 h post-inoculation), at which point they were euthanized by storage at −80°C. Each squid was homogenized and plated on LBS-Xgal, and the blue/white colony ratios were used to score these competitions as described previously (23, 55).

## Data analysis and graphing

Data analysis was conducted using Python, including the pandas library. For fluorescence of colonies, aggregates, and crypts, the mean GFP and mCherry for the region of interest and a nearby background region was acquired using Zen Blue Software. The background for each channel was subtracted from the region of interest. To normalize GFP to plasmid copy number, GFP was divided by mCherry. This resulted in the reported mean GFP/mCherry reading for each individual colony, aggregate, or crypt space. Graphpad Prism was used to construct graphs and perform statistical analyses.

## ACKNOWLEDGMENTS

We thank John F. Brooks II for contributing strains for this study and Karen L. Visick for helpful comments on the manuscript. This work was funded by NIGMS grant R35 GM119627 to M.J.M. Support for trainees was provided by NIGMS T32 GM008061 (D.A.L), T32 GM008349 (K.M.B.), and an NSF Graduate Research Fellowship to K.M.B.

