## Supplementary material for "Hybrid histidine kinase BinK represses *Vibrio fischeri* biofilm signaling at multiple developmental stages": Figures S1, S2

**SUPPLEMENTAL FIGURES**

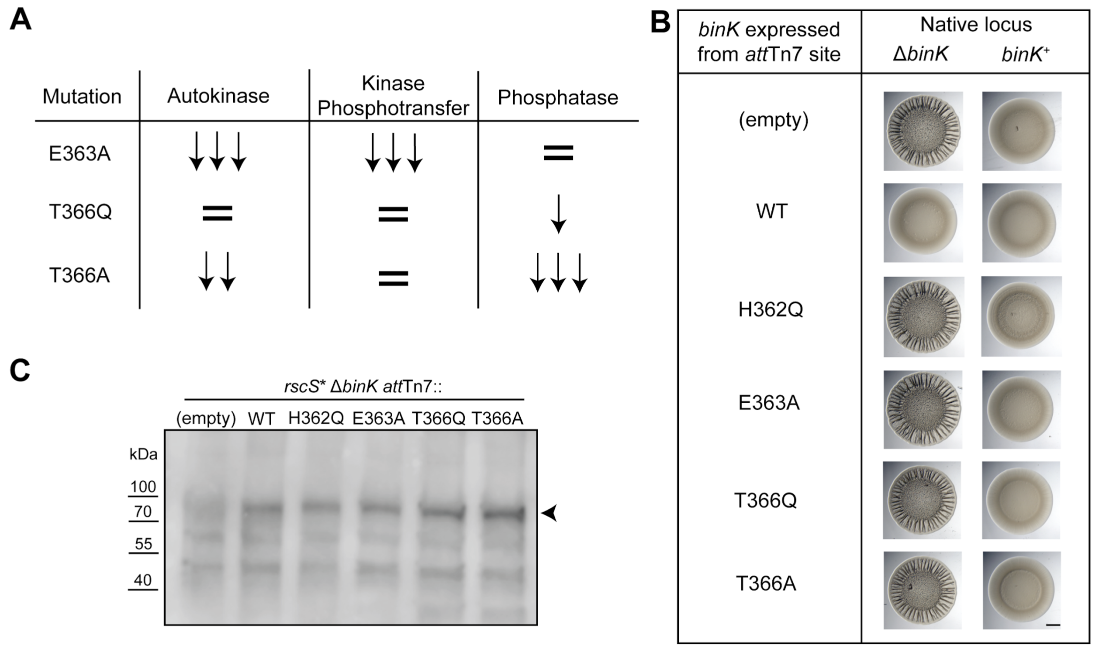


**Figure S1: Mutations in the BinK H-box region render BinK nonfunctional**

**A)** Predicted activities of H-box mutants. **B)** Wrinkled colony assay of the strains indicated, grown at 28ºC for 48 hours. In the left column, the expressed allele is the only *binK* allele in the cell, while in the right column wild-type *binK* is additionally present at its native locus. Scale bar is 2 mm. **C)** Western blot analysis of whole cell lysates assessed with the peptide antibody against BinK (arrowhead).

**Hybrid histidine kinase BinK represses *Vibrio fischeri* biofilm signaling at multiple developmental stages** - Denise A. Ludvik, Katherine M. Bultman, and Mark J. Mandel

**SUPPLEMENTAL FIGURES**

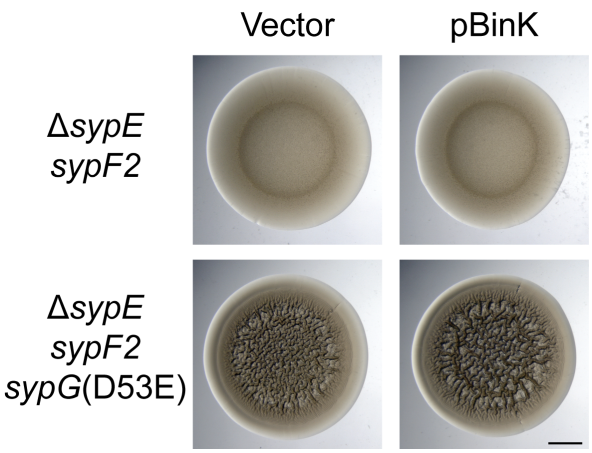


**Figure S2: Single-copy phosphomimetic SypG is epistatic to BinK overexpression**. SypG(D53E) expressed from its native chromosomal locus continues to exhibit the wrinkled colony phenotype in the Δ*sypE sypF2* background, even upon overexpression of BinK. Wrinkled colony assay on LBS agar at 25°C for 72 h. Vector is pVSV104. Scale bar is 2 mm.
